# Genomic sequences and annotations of two pseudomonas species isolated from marine and terrestrial habitats

**DOI:** 10.1101/2024.06.27.600566

**Authors:** Eric Manirakiza, Timothée Chaumier, Leïla Tirichine

**Affiliations:** Nantes Université, CNRS, US2B, UMR 6286, F-44000 Nantes, France; Institute for Marine and Antarctic Studies (IMAS), Ecology and Biodiversity Centre, University of Tasmania, Hobart, TAS, 7004, Australia

**Keywords:** *Pseudomonas stutzeri*, *Pseudomonas chlororaphis*, environmental microbiology, bacterial genomics

## Abstract

Here, we present the complete genome sequences and annotations of two species of the *Pseudomonas* genus isolated from marine and terrestrial environments. Both genomes and their annotations are available on BacBrowse (https://BacBrowse.univ-nantes.fr). This study will contribute to better understanding of the diversity present within the *Pseudomonas* genus.

---

*Pseudomonas* is one the most diverse bacterial genera, widely distributed in nature and is the genus with the highest number of species among the gram negative bacteria (1). Members of this genus show remarkable metabolic and physiological flexibility, enabling them to thrive in diverse terrestrial and aquatic environments. They are of great interest because of their importance in both plant and human diseases as well as their expanding potential in biotechnological applications (2-4). The *Pseudomonas* species used in this study are *P. chlororaphis* (NCIMB_15121) isolated from soil in Denmark and *P. stutzeri* (NCIMB_885), isolated from the Clyde Sea. Upon isolation, *P. stutzeri* was plated on nutrient agar, and *P. chlororaphis* was plated on Luria-Bertani (LB) medium (5). Both strains were retrieved from - 80°C frozen stocks, plated on solid LB medium, and individual colonies were selected for culturing in liquid LB at 30°C with shaking at 180 rpm for 48 hours. After cell harvest, genomic DNA was extracted using the Promega Wizard® HMW DNA Extraction Kit (A2920) according to manufacturer’s instructions. The DNA libraries were prepared using the SMRTbell Express Template Prep Kit 2.0 following the manufacturer’s instructions. A total of 4 μg of genomic DNA was sheared with Covaris g-TUBES at 4000 RPM. DNA was sequenced by Genome Quebec using the Pacific Biosciences Sequel platform with Circular Consensus Sequencing (CCS). CCS reads were generated from subreads with CCS 6.4.0 and assembled with Canu (v2.2) (6). Completeness and contamination levels of the assemblies were estimated using CheckM (v1.2.1) with the bacterial lineage workflow (7). The assembled genomes were then annotated using anvi’o v7.1 (8), which runs NCBI’s Clusters of Orthologous Groups database (COG20) to assign functions to genes. This revealed 5923 and 4295 coding regions for *P. chlororaphis* and *P. stutzeri* respectively. Additional gene annotations were conducted with DFAST v1.2.20 (https://dfast.ddbj.nig.ac.jp/dfc/), Prokka with compliant parameter (v1.14.6, https://github.com/tseemann/prokka) EggNOG (v2.1.12, https://github.com/eggnogdb/eggnog-mapper/wiki/eggNOG-mapper-v2.1.5-to-v2.1.12mappers). The parameters -m [diamond] and pfam_realign [realign] were used for EggNOG. Additional genome features were identified and annotated including tRNAs using tRNAscan-SE (v2.0.11) (9), tandem repeats with Tandem Repeats Finder (v4.09.1) (10) and pseudogenes were predicted by Pseudofinder (v1.1.0) (11). Genome features for both strains are summarized in Table 1.

**Table 1.** Genome features of *P. chlororaphis* and *P. stutzeri*.

|  | *P. chlororaphis* | *P. stutzeri* |
| --- | --- | --- |
| Number of CCS reads | 49,566 | 45,968 |
| Average length of CCS reads (bp) | 6979.45 | 7087.93 |
| Number of contigs | 1 | 1 |
| Mean genome coverage | 51.5X | 69.8 X |
| Genome size (bp) | 6,675,668 | 4,639,068 |
| Contamination | 0 | 0 |
| Completeness (%) | 98.28 | 100 |
| GC content (%) | 63 | 61.3 |
| Coding Region (%) | 88.3 | 89.5 |
| Average length CR (bp) | 995.51 | 966.71 |
| IR (%) | 11.9 | 10.2 |
| Average length IR (bp) | 164.90 | 147.05 |
| Total repeats | 461 | 218 |
| Total tRNAs | 73 | 63 |
| Total rRNAs | 16 | 12 |
| Total pseudogenes | 793 | 358 |

Using AntiSMASH (7.1.0) (12), the prediction of secondary metabolites revealed shared biosynthetic gene clusters (BGCs) for NRPS, RiPP, Redox cofactor, NAGGN, and Betalactone in both species. Additionally, specific BGCs including hydrogen cyanide, hserlactone, NI-siderophore, and arylpolyene were exclusively identified in *P. chlororaphis*, whereas *P. stutzeri* specifically contained BGCs for terpene production. These differences in BGCs could be attributed to their contrasting habitats, terrestrial versus aquatic, which may reflect differences in environmental conditions and microbial competitors. Both genomes are accessible in BacBrowse (https://BacBrowse.univ-nantes.fr), facilitating the exploration of *Pseudomonas* genomes and identification of bioactive compounds. Notably, identified genes and pathways can be genetically modified in tractable species like those investigated in this study.

Legend:

## ACKNOWLEDGMENTS

This study was supported by Connect Talent EpiAlg grant from Région Pays de la Loire and µAlgaNIF France-Japan International Research Project to LT. We thank Udita Chandola for DNA extraction. We are grateful to the Genomics Core Facility GenoA, member of Biogenouest and France Genomique and to the Bioinformatics Core Facility BiRD, member of Biogenouest and Institut Français de Bioinformatique (IFB) (ANR-11-INBS-0013) for the use of their resources and their technical support.

## AUTHOR CONTRIBUTIONS

LT conceived, designed and supervised the study. EM performed the bioinformatics analysis. TC contributed to the bioinformatics analysis. LT wrote the manuscript with input from EM and TC. All authors read and approved the manuscript.

## DATA AVAILABILITY

The genome sequences of *P. chlororaphis* and *P. stutzeri* have been deposited at the NCBI under assemblies ASM3849947v1 and ASM3850004v1, respectively (BioProject PRJNA1101843). The annotations are available on Zenodo under the following DOI: https://zenodo.org/doi/10.5281/zenodo.11657795.

## Notes

https://zenodo.org/doi/10.5281/zenodo.11657795

### Competing Interest Statement

The authors have declared no competing interest.

